# The Main Themes and Experiences Shared with the Hashtag #BlackintheIvory: A Qualitative Analysis

**DOI:** 10.1101/2022.04.13.488234

**Authors:** Mopileola Tomi Adewumi, Paul Delgado, Marvin Carr, Emily Sowah, Matt Vassar

**Author notes:** **Corresponding Author:** Mr. Marvin Carr. Address: 1111 W 17th St., Tulsa, OK 74107, United States.

## Abstract

**Background:** The movement #BlackintheIvory gave Black academics an opportunity to connect through a social media platform allowing them to share common experiences in the pursuit of higher education. Through the analysis of Twitter posts, using the hashtag #BlackintheIvory, this study investigates the main themes identified among Black scholars in academia and their shared experiences with teaching, mentoring, collegiality, identity, service, and racism.

**Methods:** Using the Twitter API, we isolated all publicly available tweets, which can include text, images, and links to websites, posted with the hashtag #BlackintheIvory on the Twitter website (www.twitter.com) from the inception of the hashtag in June 2020 to the end of December 2020. To evaluate the tweets, we categorized the tweets inductively. Based on the content of the posts, we identified 6 themes: Teaching, Mentoring, Collegiality, Identity, Service, and Racism.

**Results:** Our search yielded a total of 12,538 original posts, including tweets between inception in June 2020 to December 2020 from profiles made public (excluding modified tweets and duplicate tweets). We selected and analyzed the top retweeted 2500 tweets, which is 20% (2500/12538) of the total number of downloaded tweets. The greatest percentage of posts were about Teaching (881; 35%), followed by Service (441; 18%) and Racism (414; 17%). The remaining tweets were categorized as Collegiality (388; 15%), Identity (210; 8%), and Mentoring (166; 7%) of the total number of tweets from June-December 2020.

**Conclusion:** The experiences, perspectives, and narratives among the Black diaspora within #BlackintheIvory are not uniform. The commonality exists within the structural systemic racism which impacts Black academics within the ivory tower, this study is a resounding wake-up call for action.

**Funding:** This study was not subject to any funding

## Introduction

The disturbing event between George Floyd and the Minneapolis police department highlighted the long-standing racial inequalities and systemic problems of police violence existing in the United States (1). The repeated deaths of Black people including George Floyd, Breonna Taylor, and Ahmaud Arbery, resulted in protests across the United States and marches globally in the year 2020 (1, 4). Along with the physical demonstrations across the country, people joined openly calling for action on racism on social media (5). Over the last decade, social media outlets have become a major medium of communication and discussion, especially about controversial topics. ^1^ About 75% of the US adult population have reported using Twitter as a news outlet^2^. Hashtags have become a way to start conversations about any topic, this allows social media posts on such topics to be easily found and shared, opening discussions and allowing users to share their opinions.

The hashtag #BlackintheIvory, a social media movement started on Twitter, was, therefore, an opportunity to highlight many of the inequities experienced among Black scholars (1). Shardé Davis and Joy Melody Woods were key players in the development and creation of #BlackintheIvory, a movement resulting from a conversation between two Black scholars reflecting on their own journey in academia (7). Although the hashtag #BlackintheIvory started as an exchange of personal stories, it highlighted the common shared experiences among Black scholars.

The term ‘ivory tower’ has previously examined the complex ethical and social issues facing disproportionately modern institutions. The constant discriminatory practices experienced by underrepresented minority scholars have resulted in challenges undermining aspects of inclusion (10, 11). In 2018, of the 1.5 million faculty in degree-granting postsecondary institutions, Black males and Black females each accounted for only 2% of full-time professors (12). The movement #BlackintheIvory gave Black academics an opportunity to connect through a social media platform allowing the sharing of common experiences. Through analysis of thousands of retweets using #BlackintheIvory, this study, investigates the main themes identified among Black scholars in academia and their shared experiences with teaching, mentoring, collegiality, identity, service, and racism.

## Methods

In this study, we sought to determine the impact of the hashtag #BlackintheIvory, its reach, passed messages, and to put together an analysis of the stories and experiences shared by Black academics using the hashtag.

### Tweet Selection

Using the Twitter API, we isolated all publicly available tweets posted with #BlackintheIvory on the Twitter website (www.twitter.com), from the inception of the hashtag in June 2020 to the end of December 2020. Figure 1 demonstrates a word cloud of the top 100 hashtags shared in combination with #BlackintheIvory. The identified tweets were further stratified by sorting the tweets by retweets to capture the most retweeted tweets, to include the tweets with the most impact and most reach. We then selected and analyzed the top 20% of the most retweeted tweets.

**Figure 1:**
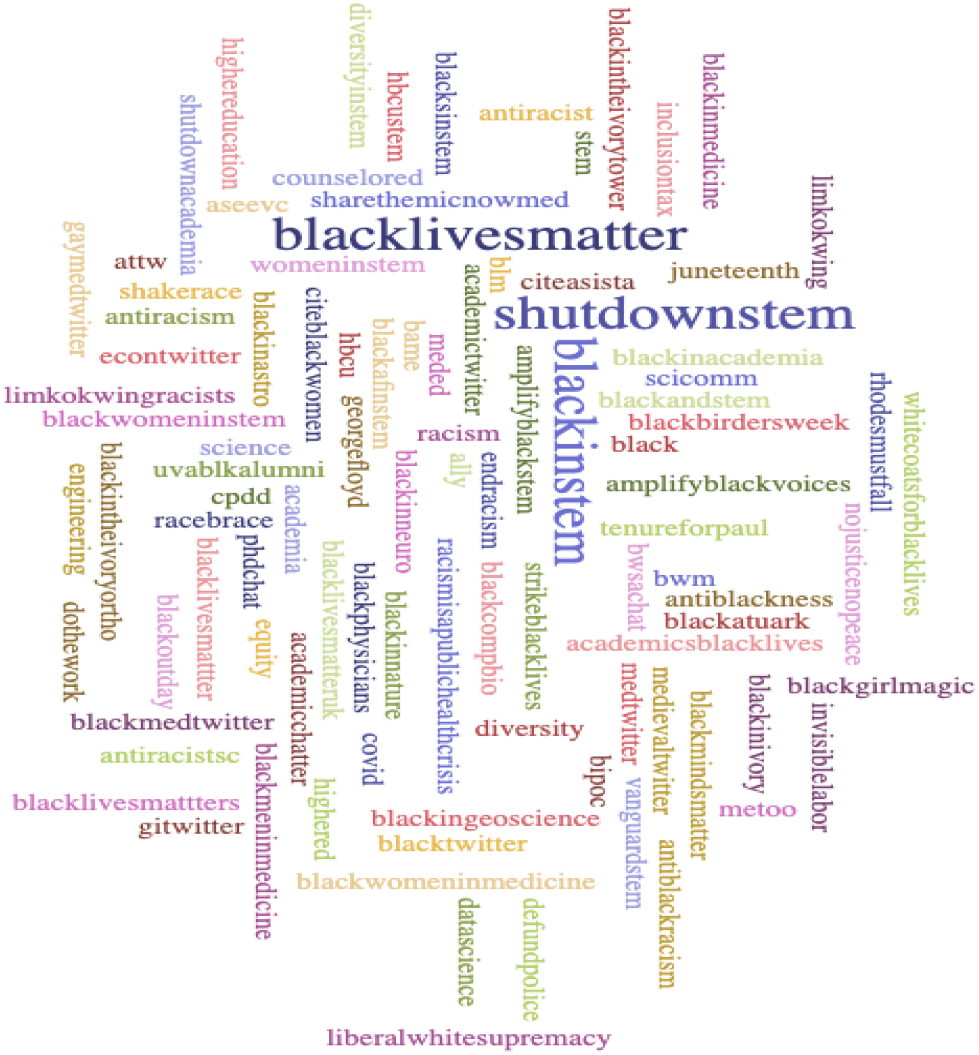
Word cloud of the top 100 hashtags shared in combination with #BlackintheIvory tweets

### Themes and Categories

To evaluate the tweets, we categorized the tweets based on the experiences shared by the users. A previous study by Stanley examined the experiences of several faculty of color working in predominantly white institutions (17). By using content and narrative analysis, Stanley grouped their stories into 6 themes: Teaching, Mentoring, Collegiality, Identity, Service, and Racism. We thus decided to group our selected tweets into categories using the themes from Stanley’s study (17), as we believe they are representative of experiences among academics. Table 1 contains the description of each one of the six categories with examples of tweets representing each.

**Table 1:**
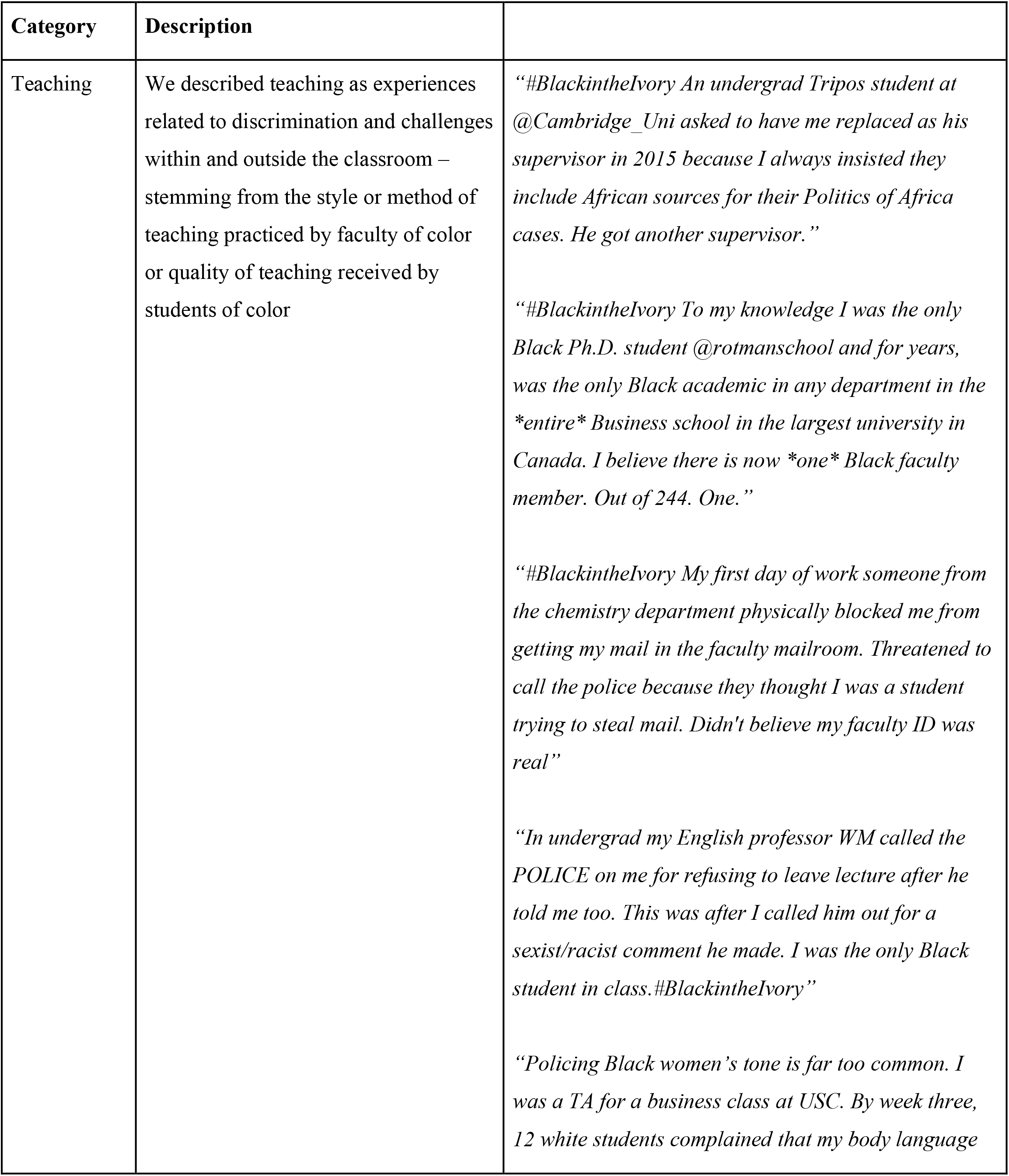
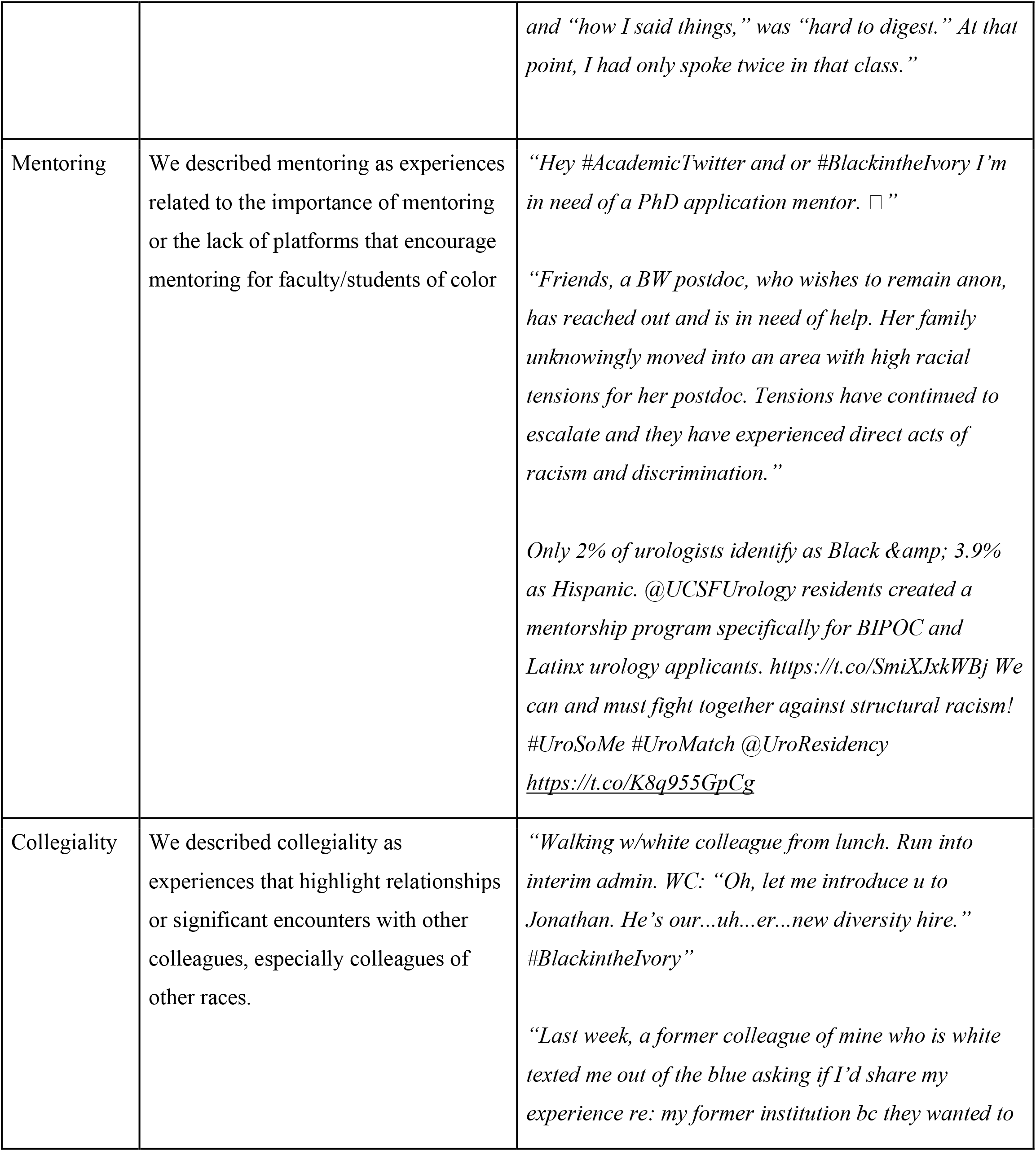
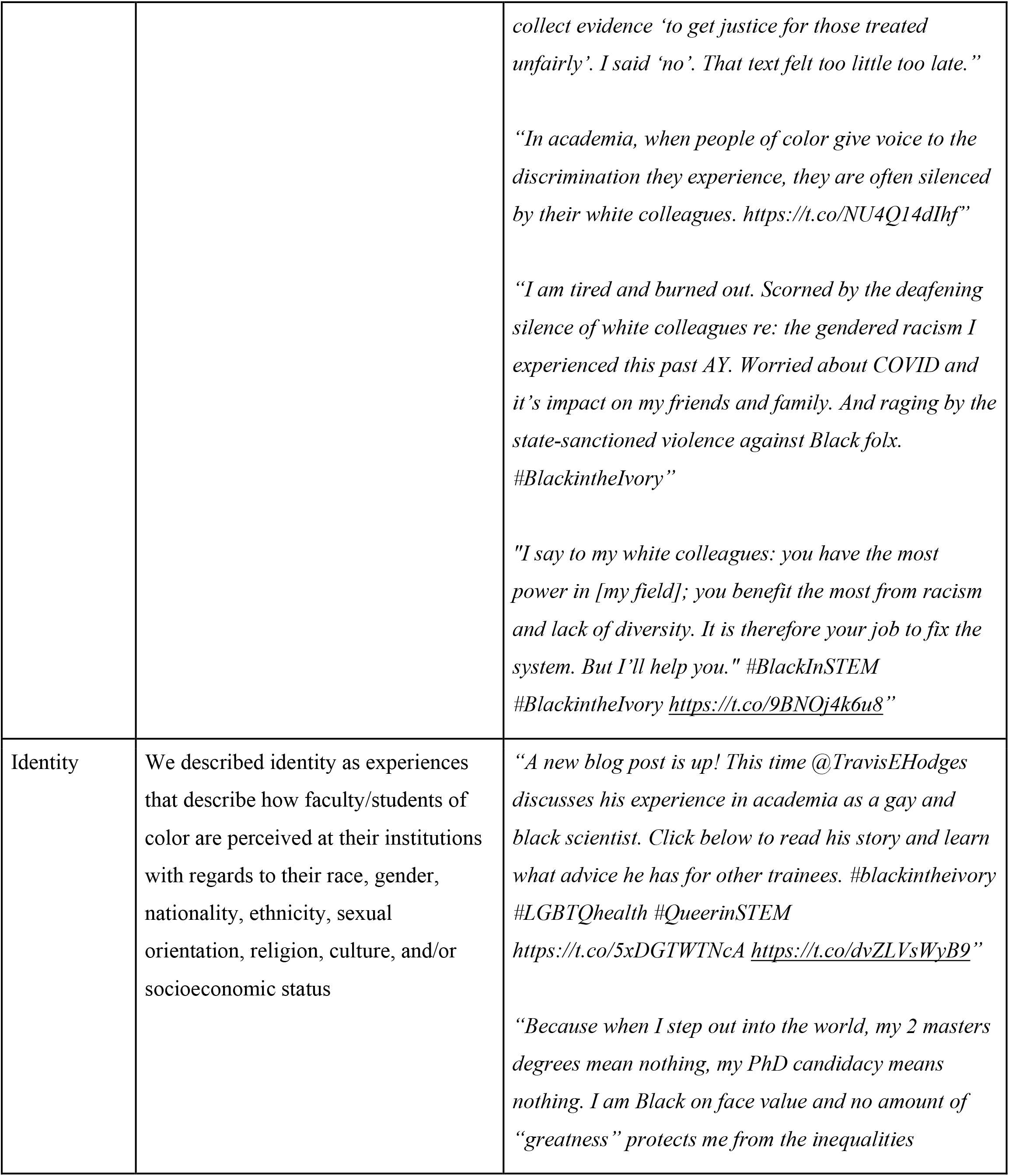
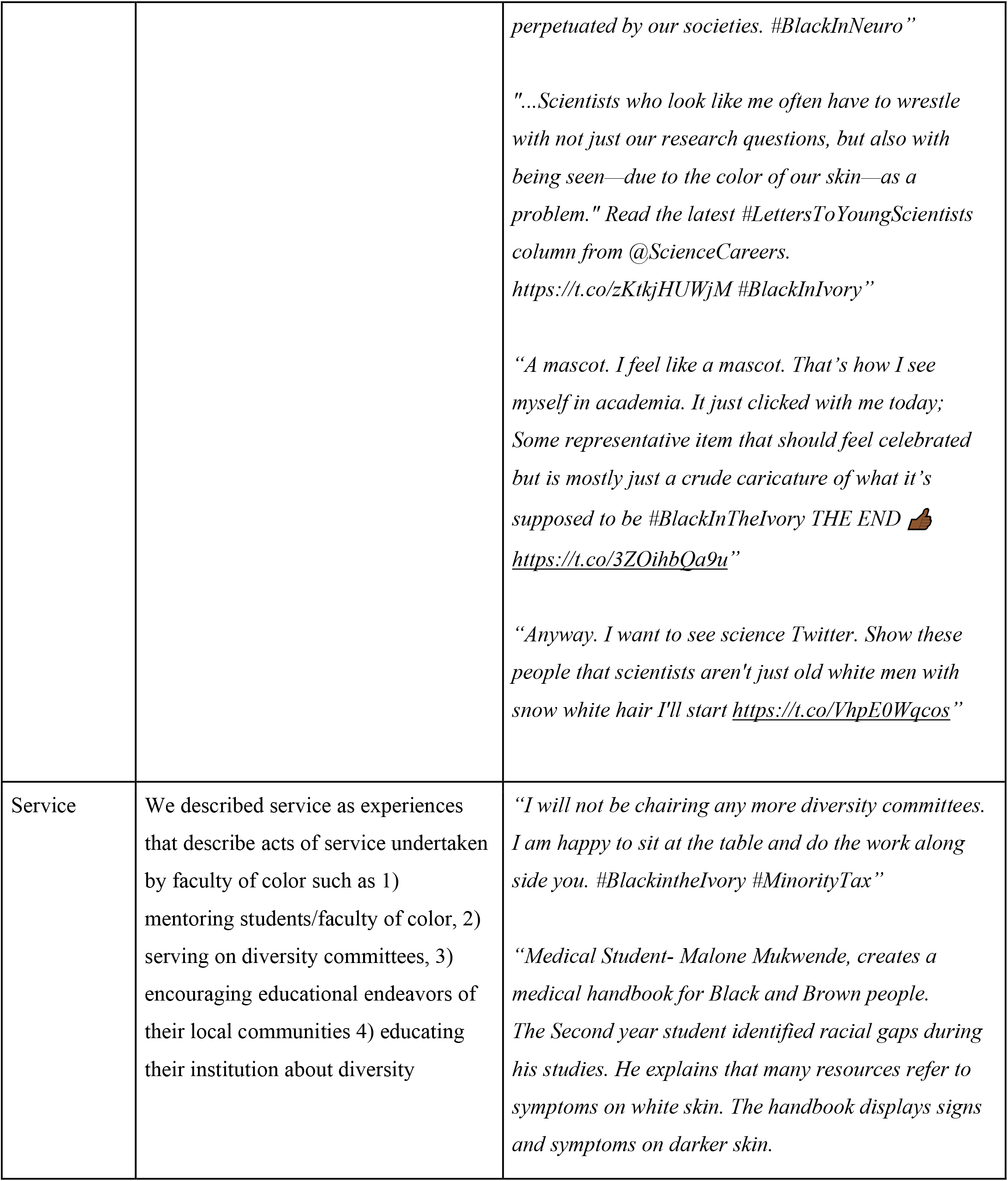
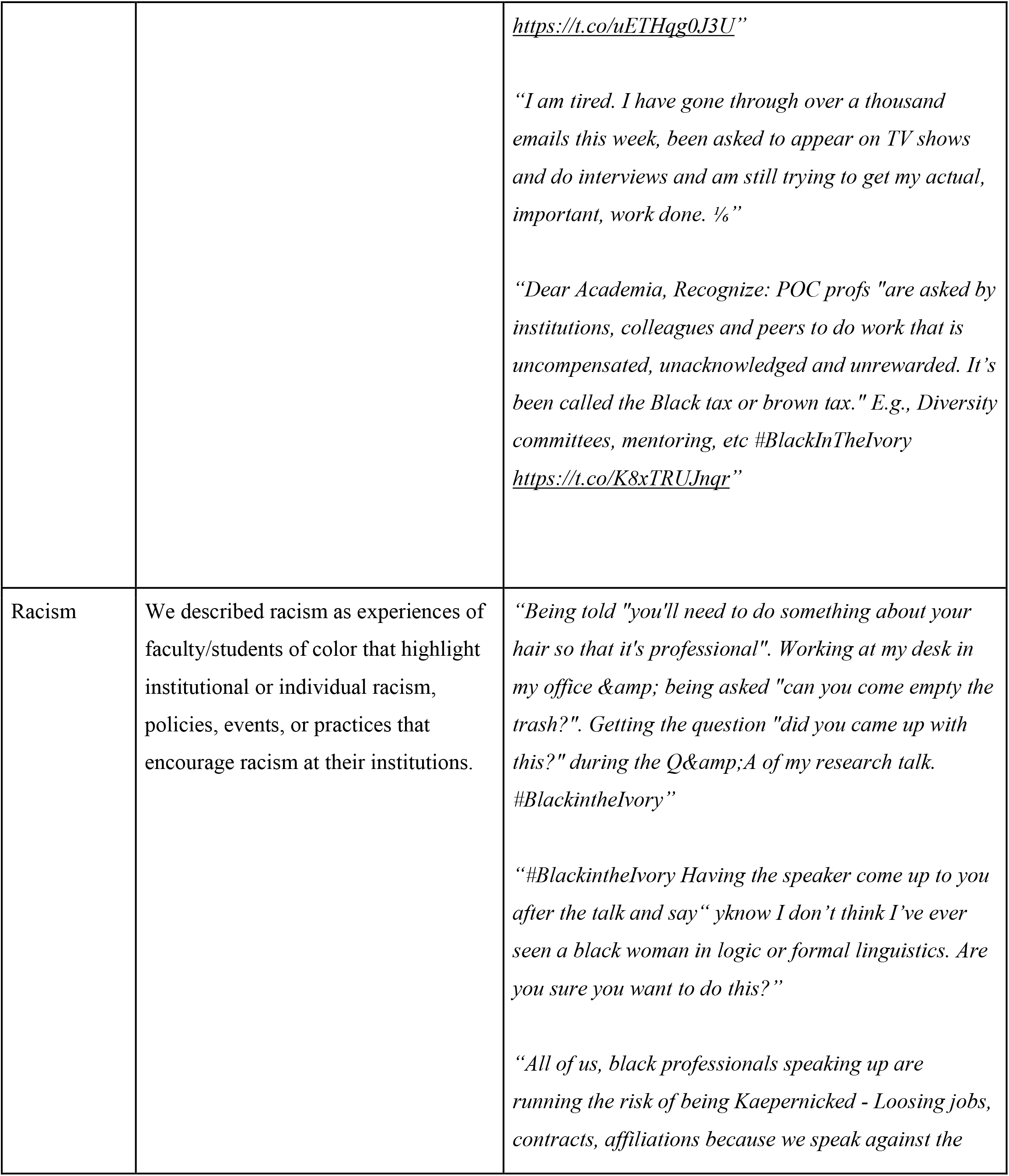
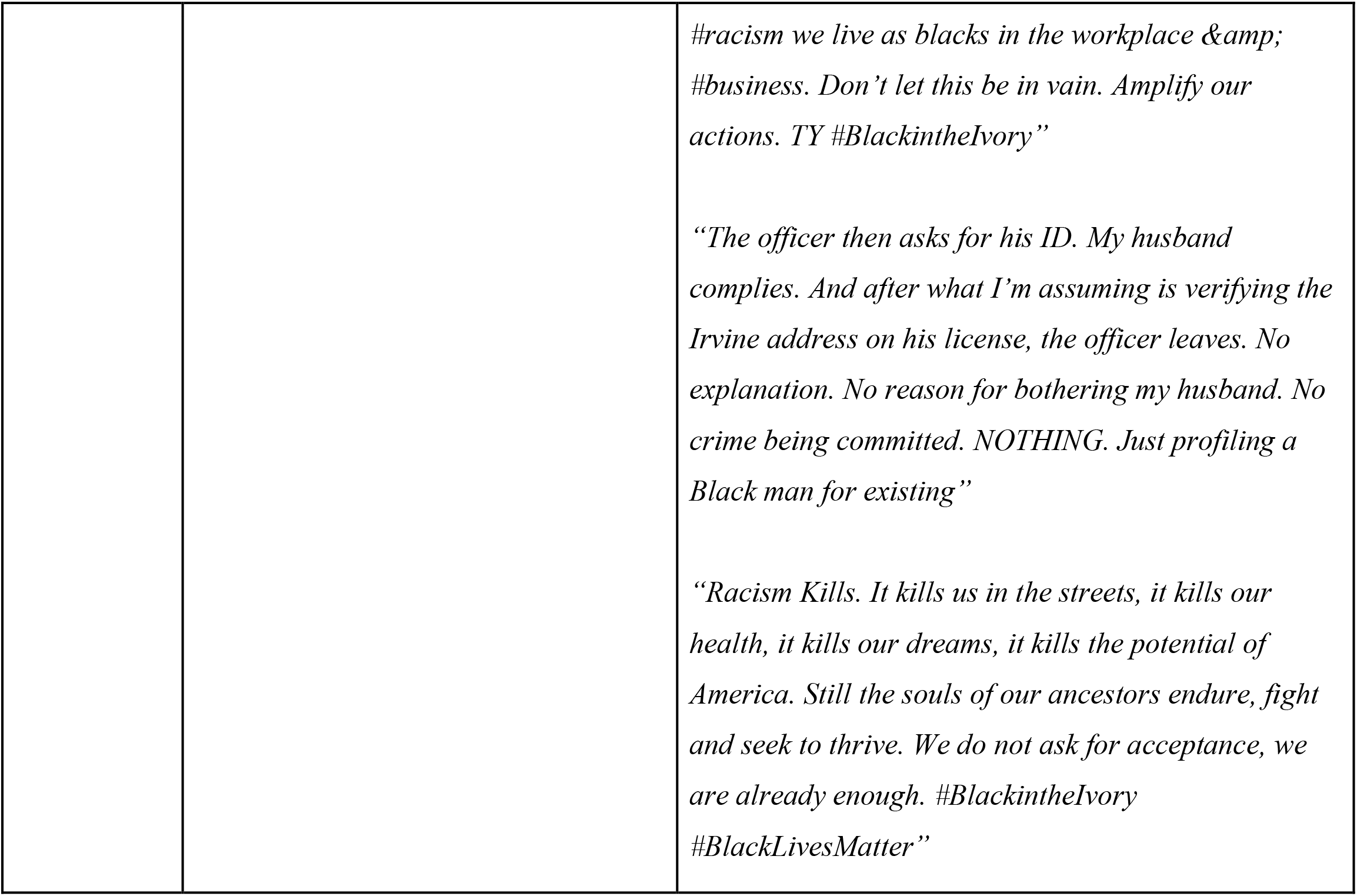
Major themes of experiences, definitions and examples identified in the analysis of the #BlackinTheivory.

### Training

To ensure consistency between investigators, TA provided in-person training to PD and ES prior to data extraction. During this training session, we (1) described the aim and objectives of the study, (2) reviewed the study protocol, (3) described and shared our perspectives on each theme being assessed (4) discussed how to extract the said data points and match tweets with selected themes.

### Data Extraction and Analysis

After training, PD and ES extracted data in a duplicate, independent, and blinded fashion. Extraction began on January 23, 2021, and ended on March 31, 2021. Responses were discussed between investigators and discrepancies were resolved by TA if needed.

To evaluate the experiences expressed by the users of #BlackintheIvory, we used a grounded theory and phenomenological research method to analyze the tweets. We approached each of the 2500 selected tweets with open-coding, memo-writing, and categorization to determine and critically understand the narratives and experiences being shared in each tweet.

## Results

### General Impact

Our search yielded a total of 12,538 original tweets between inception in June 2020 to December 2020 from public profiles (excluding retweets, modified tweets, and duplicate tweets). We selected and analyzed the top retweeted 2500 tweets, which is 20% (2500/12538) of the total number of downloaded tweets while accounting for the volume of tweets from month to month. The majority (64%) of the tweets were shared in the month of June, which was the month following the police shooting of George Floyd, with protests erupting around the country. Many users took the platform to express their reflections of what it means to be Black in academia. Table 2 & 3 contains the different forms of involvement among Twitter users and the #BlackintheIvory movement (i.e., tweets, likes, replies, retweets). Between June and December 2020, a total of 2,608,029 retweets and 9,722,382 likes were shared by Twitter users using the #BlackintheIvory. These tweets also led to continued conversations with a total of 238,896 unique replies shared by users using #BlackintheIvory across Twitter.

**Table 2:** Total Number of tweets vs Number of Tweets selected for analysis per month.

| **Month** | **Number of Tweets** | **Number Tweets Selected for Analysis (% of total tweets)** |
| --- | --- | --- |
| ***June*** | 7996 (64%) | 1600 (12.8%) |
| ***July*** | 1939 (16%) | 400 (3.2%) |
| ***August*** | 1042 (8%) | 200 (1.6%) |
| ***September*** | 579 (5%) | 125 (1%) |
| ***October*** | 552 (4%) | 100 (0.8%) |
| ***November*** | 311 (2%) | 50 (0.4%) |
| ***December*** | 119 (1%) | 25 (0.2%) |
| **Total** | **12538 (100%)** | **2500 (20%)** |

**Table 3:** Total Number of Tweets per Month.

| **Month** | **Number of**<br>**Tweets** | **Number of**<br>**Likes** | **Number of**<br>**Replies** | **Number of**<br>**Retweets** |
| --- | --- | --- | --- | --- |
| ***June*** | 7996 (64%) | 7216203 | 161747 | 1855483 |
| ***July*** | 1939 (16%) | 1655799 | 26637 | 512954 |
| ***August*** | 1042 (8%) | 475023 | 13220 | 133752 |
| ***September*** | 579 (5%) | 198777 | 20793 | 64244 |
| ***October*** | 552 (4%) | 99878 | 13194 | 25360 |
| ***November*** | 311 (2%) | 57311 | 2679 | 10204 |
| ***December*** | 119 (1%) | 19391 | 626 | 6032 |
| **Total** | **12538 (100%)** | **9.722,382** | **238,896** | **2,608,029** |

### Themes

As shown in Table 4, the greatest percentage of posts were about Teaching (881; 35%), followed by Service (441; 18%), then followed closely by Racism (414; 17%). The remaining tweets were categorized as Collegiality (388; 15%), Identity (210; 8%), and Mentoring (166; 7%) of the total number of tweets from the months June-December 2020.

**Table 4:**
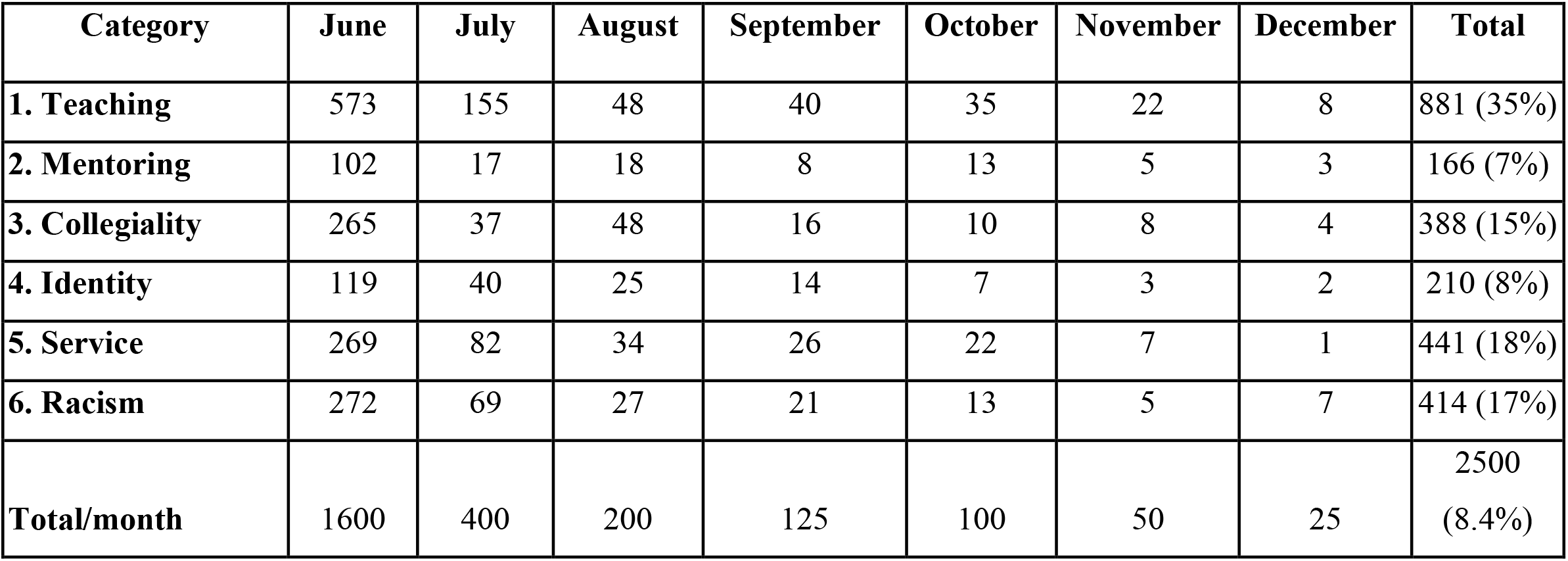
Number of tweets from months June-December of 2020 based on the six main theme categories.

| Category | June | July | August | September | October | November | December | Total |
| --- | --- | --- | --- | --- | --- | --- | --- | --- |
| **1. Teaching** | 573 | 155 | 48 | 40 | 35 | 22 | 8 | 881 (35%) |
| **2. Mentoring** | 102 | 17 | 18 | 8 | 13 | 5 | 3 | 166 (7%) |
| **3. Collegiality** | 265 | 37 | 48 | 16 | 10 | 8 | 4 | 388 (15%) |
| **4. Identity** | 119 | 40 | 25 | 14 | 7 | 3 | 2 | 210 (8%) |
| **5. Service** | 269 | 82 | 34 | 26 | 22 | 7 | 1 | 441 (18%) |
| **6. Racism** | 272 | 69 | 27 | 21 | 13 | 5 | 7 | 414 (17%) |
| **Total/month** | 1600 | 400 | 200 | 125 | 100 | 50 | 25 | 2500<br>(8.4%) |

#### 1. Teaching

The most discussed theme within our dataset was the teaching experience of Black academia (881; 35%). Many posts described the discrimination and microaggressions faced by Black academics, most posts were about the challenges and discrimination they faced from their students — how they are perceived by their students, fellow faculty, or their academic institutions.

> *“Recent student evaluation stated that I should never teach again & I had committed malpractice since I made race central to American Hist. & My worst stated that I should be ashamed that I was pregnant &; Black w/ no husband as I was a walking stereotype– #BlackintheIvory“*

Many users also took the time to point out issues arisen as they try to incorporate issues about diversity or inclusion in their course content.

> *“When I applied for teaching positions, I was encouraged to remove my social justice work from my resume and anything “too pro-black” from my cover letters. I was told to decide if I wanted to be a great teacher in general or a great teacher for black students.” #BlackintheIvory*

Many tweets also came from students expressing their discontent with the amount of diversity in their classes, their faculty choices, and the lack of representation in the courses being taught.

> *“Having to take a class in the Africana Studies department just so that I could experience one black Professor during my Masters experience #BlackintheIvory”*

Overall, academic centers are specifically created to allow for teaching and these tweets show how race plays a part in the teaching experiences of students and faculty alike in these academic institutions.

#### 2. Service

The second most discussed theme in our dataset was service. many users described services they have had to provide or be engaged in as faculty or students of color at their different institutions. Some express the expectation coming with their position in academia such as mentoring other underrepresented colleagues, educating their colleagues on issues of diversity, serving on numerous diversity committees, and giving back to their communities.

> *“I’m not waiting for universities to do diversity & inclusion right. I’m launching something w/ my own resources to help many #BlackintheIvory folks. Unis don’t work fast enough. Diversity officers have no power. People in the academy are too afraid. Stay tuned.”*

Some tweets also used the platform to recognize and share several services being provided by minority academics across different fields.

> *“Today, our student-athletes launch #DamChange, a platform to educate, empower, and enhance the experience of Black student-athletes and staff. Take two minutes. Hear their experiences. Then help us create #DamChange. Learn More: https://t.co/67o5Ulumac#GoBeavs”*

Although faculty and students alike express the willingness to provide these services, many users used the platform to also shed light on the burden and extra stress such service places on them — in addition to the regular work expected of them, be it scholarly work, studying or administrative work. This concept was described by many as *“minority tax”* which is defined as “the burden of extra responsibilities placed on minority faculty in the name of diversity” ^18^

> *“This week alone, I have spent over 14 hours in meetings discussing Diversity and Inclusion efforts at our med school. While important work, I could have used this time for schoolwork and research projects. This is what the minority tax looks like. #BlackintheIvory“*

One of the major issues making service cumbersome and burdensome for several academics is the lack of compensation or recognition for the work being done.

> *“#BlackInTheIvory damned if you do, damned if you don’t get involved with diversity and inclusion work. If you don’t, you know it’ll end up being a white people’s version of diversity, if you do, you end up using research time for work that won’t even be recognized”*

#### 3. Racism

Racism was unsurprisingly one of the most discussed topic categories (414; 17%). Many users took the opportunity to describe experiences highlighting microaggression, discrimination, stereotype, or disadvantages they have faced based on their ethnicity, race, nationality, or background.

> *“What are some #microaggressions you have received as a person of color in academia? Here are some I have received: 1) you speak so well were you adopted 2) you talk so well did your read a lot as a kid 3) your error bars are too small do you know how to do stats”*

Many users also took time to point out the effects these experiences have had on their productivity at work and well-being outside of work.

> *“The racism Black women endure in academia & medicine is a public health crisis. Our days are spent in spaces that attempt to restrict our voices, invalidate our work: limit our impact. This stress heightens our risk of physical, reproductive & mental health disorders. A thread”*

The fear of retribution is a factor preventing many academics of color from speaking up about racist experiences.

> *“Instead of acting on this, apologizing, supporting me, or even acknowledging harm, I was lectured, silenced, and sidelined for writing this tweet. I’m still being punished for publicly naming racism. And I will be punished for writing this as well. #BlackintheIvory”*

Lastly, the platform allowed for some people to share ideas about moving forward and the steps we need to take as a community to solve racism.

> *“Dismantling structural racism in medicine is a collective responsibility, and everyone has a role”*

#### 4. Collegiality

Many users tweeted about encounters and relationships with their colleagues in academia shaping their experiences as academics of color at their various institutions.

> *“At a professional meeting. My friend said to his colleague, “I’d like to introduce you to the next @ACSMNews president,” gestured toward me. As I held out my hand to greet him, his colleague turned, smiled excitedly, looked past me, and said, “Where is he?”#BlackintheIvory”*

As academics of color go through their everyday struggles, they sometimes look to others (their colleagues) for support. Many users used the platform to highlight — the lack of awareness or action on the part of their white colleagues, in supporting their struggles as people of color.

> *“I discuss systemic racism in all my classes. Constantly I am told I center race too much. And now all my white colleagues wanna use me to write statements because they just discovered what systemic racism is watching the passing of George Floyd 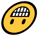 #blackintheivory”*

Users went on to describe and highlight the roles their colleagues can play in fixing these issues, emphasizing their white colleagues possess the ability to help even when they do not realize it.

> *“A lot of non-Black friends and colleagues have asked what they could do today for #Strike4BlackLives. Since you likely know who your Black colleagues are talking about in #BlackintheIvory, one could start by confronting the racism they describe from folks in your department”*

Some also used the opportunity to reach out to other colleagues of color for support and ideas.

> *“Dear #BlackInTheIvoryTower colleagues and friends, I see you, I hear you, and most importantly, I believe you. For my small part today, I pledge to re-examine and update my syllabi. #AcademicTwitter, join me. Welcoming medical sociology suggestions”*

#### 5. Identity

People identify themselves differently, our identification with certain nationalities, races, religions, or sexual orientation. Interestingly, many users of the hashtag talked about experiences and interactions stemming from how they choose to identify themselves and the effect of these experiences on them.

> *“As a woman of color in academia with natural hair, I would love to get to the point when I don’t worry about my hair being perceived as unprofessional or a ploy to exclude me from opportunities. Oh well, I’m not changing my look! @AcademicChatter @TweeterAcademic @BlackWomenPhDs”*

Many users also used the opportunity to express how their identities have been exploited or misused by people and by their institutions. Using the opportunity to enlighten others about these issues.

> *“PSA: Do not use ‘Black’ as a noun to describe a person or group of people (ex. ‘a Black’ or ‘Blacks’ is a no-no and further it is dehumanizing). Use ‘Black’ as an adjective only (ex. Black woman, Black people.’ And don’t forget upper-case ‘B’.”*

Lack of support and a sense of community plays a huge role in the effects these encounters have on people of color. Many users also used the opportunity to preach acceptance, and encourage others to be proud of their identity.

> *“This is me. My authentic self. It took me years to find my voice. But now I have. And I will continue to speak. Because these are the conversations we need to have. I am first, and foremost, a black woman. I love the skin I’m in. #BlackLivesMatter #BlackintheIvory”*

#### 6. Mentoring

Users described the importance of mentorship and the role mentors have played in their lives. Many users used the hashtag as an opportunity to celebrate achievements made possible through mentorship.

> Thank you, everyone, for all the support, it took a village to get me here and I am grateful to all my mentors, friends and family. Successfully defended my PhD. I’m a PhD!!!!
>
> #blackimmunologist #BlackGirlMagic #BlackintheIvory #blackinstem

Mentorship works differently for underrepresented scholars, especially in academia, mostly due to the lack of representation within the system. Several users expressed their frustration and the difficulty of finding mentors of color.

> *“So I pulled the data on race/ethnicity of tenure line Dartmouth faculty. There are 14 Black scholars on a faculty of 459 people (3.1%). This means we have a lower % of Black faculty today than we did in 2004. I know our peers aren’t much better, but the #’s still make me angry”*

Many academics of color fall back on platforms such as Twitter to find mentors, and many used the #blackintheivory as a platform to share programs supporting this.

> *“We are thrilled to introduce the @PGSFellowship! This fellowship aims to empower Black excellence in aerospace by providing extraordinary Black undergrads with paid summer internships as well as executive and peer mentorship! Read the official announcement here “*

## Discussion

The use of #BlackintheIvory has become a national soundboard on Twitter for Black academics cohabitating within the ‘ivory tower’. This state of seclusion from the practicalities of the real world by the ‘ivory tower’ has uprooted civil unrest. At the age of six Ruby Bridges became the first Black scholar to integrate into an all-white school in the south. At the tail end of the Jim Crow era, Ruby defied the odds of blatant racism and bigotry becoming one who didn’t follow the path yet blazed a new trial. In her book titled “Through My Eyes”, Ruby states “Racism is a grown-up disease, we must stop using our children to spread it” (20). As with Ruby trailblazing the path of educational advancement for the Black community, so are the 5% of Black academics within their respective predominantly white institutions (PWI) (21).

### The Black Experience

Many Black academics using the #BlackintheIvory voiced the extra burden of taking on additional diversity commitments of their institutions minus compensation. This paradox of culture and support among Blacks within academia on the pursuit of tenure represents “Cultural taxation”, a school of thought originally studied by Padilla (1994) (22). Blacks are inherently aware of this increase in responsibilities and expectations on their climb up the educational ladder (23). Tenure is a lifelong appointment with increased compensation and in 2018 only 5% of Black were tenured (24,25). Over 70% of Black professors reported: “feeling a need to work harder than their colleagues to be seen as legitimate scholars” compared to less than half of white professors (26). In the pursuit of tenured seeking Black professors the increased work of diversity may seem insignificant or less valued to other colleagues or a school’s board of trustees.

### Strength & Limitations

Although Twitter is commonly accepted and widely used in the scientific community, the usage of this tool should be viewed considering its limitations. Although our selected keywords were among the most popular search inquiries, we recognize there may be other keywords capable of assessing public interest not included in our sample. We recognize Twitter may not display a complete representation of public interest in a specific topic thus our results may be subject to sampling bias.

## Conclusion

The experiences, perspectives, and narratives among the Black diaspora within #BlackintheIvory are not uniform. The commonality exists within the structural systemic racism which impacts Black academics within the ivory tower, this study is a resounding wake-up call for action for us all.

## Additional Contributions

We thank Dr. Urmimala Sarkar, MD., MPH for her guidance in the development, design and vetting of this study.

